# A multi-state dynamic process confers mechano-adaptation to the bacterial flagellar motor

**DOI:** 10.1101/2021.12.22.473861

**Authors:** Navish Wadhwa, Alberto Sassi, Howard C. Berg, Yuhai Tu

## Abstract

Adaptation is a defining feature of living systems. The bacterial flagellar motor adapts to changes in the external mechanical environment by adding or removing torque-generating stator units. However, the molecular mechanism for mechanosensitive motor remodeling remains unclear. Here, we induced stator disassembly using electrorotation, followed by the time-dependent assembly of the individual stator units into the motor after electrorotation was terminated. From these experiments, we extracted detailed statistics of the dwell times that comprise the stochastic dynamics of the binding and unbinding of stator units. Dwell times reveal multiple timescales, indicating the existence of multiple binding states of the stator units. Based on these results, we propose a minimal model in which the stator unit can occupy four different states – two bound states with very different rates of unbinding, a diffusive unbound state, and a transiently detached state. Our minimal model quantitatively explains multiple features of the experimental data and allows us to determine the transition rates among all four states. Our experiments and modeling suggest a mechanism of mechano-adaptive remodeling of the bacterial flagellar motor in which torque generated by bound stator units controls their effective unbinding rate by modulating the transition between the two bound states. Furthermore, the binding rate of stator units with the motor has a non-monotonic dependence on the number of bound units, likely because of two counteracting effects of motor rotation on the binding process.

---

Through billions of years of evolution, bacteria have devised myriad ways to move around^1^. One of the most common is swimming in aqueous environments by the rotation of thin helical filaments called flagella^2,3^. Flagellar rotation is powered by a biological nanomachine, the flagellar motor. This motor has been the subject of numerous studies and is often considered a model macromolecular machine^3,4^.

The structure and composition of the flagellar motor are largely conserved across bacterial species^5^. It consists of a rotor that rotates with respect to the cell, a stator that remains stationary with respect to the cell, and a rigid rod (drive shaft) and a flexible hook (universal coupling) that transmit the rotation to the filament^6,7^. The stator itself consists of individual units, each of which is embedded in the inner membrane and generates part of the torque required to drive the motor^8^. Each stator unit contains two ion channels (proton channels in *Escherichia coli*), and the passage of ions into the cell through these channels in response to the ion motive force is believed to generate rotation within the stator unit^9–11^. On the periplasmic side, the stator unit binds to the peptidoglycan layer of the cell wall, while the cytoplasmic end interacts with the cytoplasmic portion of the rotor (c-ring) to apply a torque on the rotor^8,12,13^. Furthermore, the stator undergoes dynamic turnover; bound stator units are released, and free units from the inner membrane-embedded pool bind at the periphery of the motor and increase torque production^14,15^.

A central feature of living systems is their ability to adapt to changes in their environment. How macromolecular complexes adapt remains an open question. The flagellar motor has emerged as an example of a macromolecular complex that adapts to its environment by modulating its assembly in response to chemical and mechanical signals^16,17^. For instance, in response to changes in the external mechanical load, the stator remodels by adding or removing individual stator units^18,19^. At high load, the motor is driven by a larger number of stator units, which increases its torque output^20–22^. Conversely, at low load, the motor is driven by fewer stator units, which decreases torque output and conserves energy^20,21,23^. Stator remodeling enables rapid and continuous mechano-adaptation of the motor to its external environment^17^.

Previous work modeled the assembly of a stator unit into the motor as a two-state process, with a bound (“on”) and an unbound (“off”) state. The rate of transition from the bound to unbound state decreases with increasing motor torque in response to higher load^20–22,24^. While this simple model captures population-averaged kinetics, its broader validity has not been thoroughly tested. Indeed, by analyzing the steady state distribution of the time intervals between binding and unbinding events in single motors, Shi *et al*. discovered the existence of an additional short-lived “hidden” state in which the stator unit transiently disengages from the rotor while remaining assembled in the motor^25^. This added complexity suggests that, to gain a deeper understanding of the mechanisms that enable adaptive stator remodeling, we must go beyond population-averaged data and analyze the statistics of single binding and unbinding events in individual flagellar motors. The motor behavior must be described not just in steady state but also during the adaptation process.

Here, we present a detailed analysis of remodeling events in single flagellar motors as they adapt to a sudden increase in load. From the distribution of dwell times of these events, we find signatures of additional states beyond the three (bound, unbound, hidden) that have been previously described. Motivated by these observations, we propose a new model with four states (unbound, loosely bound, tightly bound, and hidden). This treatment provides a coarse-grained approximation for a more complete model in which a bound stator unit can occupy a continuum of states with different dissociation rates. We present a theoretical treatment of the coarse-grained model using first-passage-time analysis, which allows us to calculate key statistics for the dwell time analytically. Our coarse-grained model demonstrates excellent quantitative agreement with the dwell-time statistics extracted from experimental data. Finally, we discover new features of the on-process, whereby the on-rate of an incoming stator unit is a non-monotonic function of the number of previously bound units. Put together, this work reveals that the mechanosensitive remodeling of the flagellar motor is powered by molecular interactions that are more nuanced and complex than previously thought.

## Results

### Single-motor electrorotation experiments

To measure the dynamics of flagellar motors during the adaptation process accurately, we conducted a large number of single-cell experiments with the bacterium *Escherichia coli*. We tethered individual *E. coli* cells to a surface via a short flagellar stub, causing the motor to rotate the cell body instead of the flagellar filament and thus operate under a very high load (**Fig. 1a**). In the initial phase of the experiment, we observed the cell rotation for 30 s. We then applied a high frequency rotating electric field to the cell, which exerted a large external torque on it in the same direction as the motor torque. This greatly reduced the load acting on the motor and stimulated mechano-adaptation in the motor via the release of bound stator units. We kept electrorotation ON for a total of 6 minutes. At the end of 6 minutes, we turned electrorotation OFF, which suddenly increased motor load. Once again, the motor adapted, this time by the sequential addition of stator units, with the motor speed increasing in a stepwise fashion over time (**Fig. 1b**). This phase of the experiment also lasted for 6 minutes, during which we made continuous measurements of the motor speed at a high temporal resolution.

**Figure 1.**
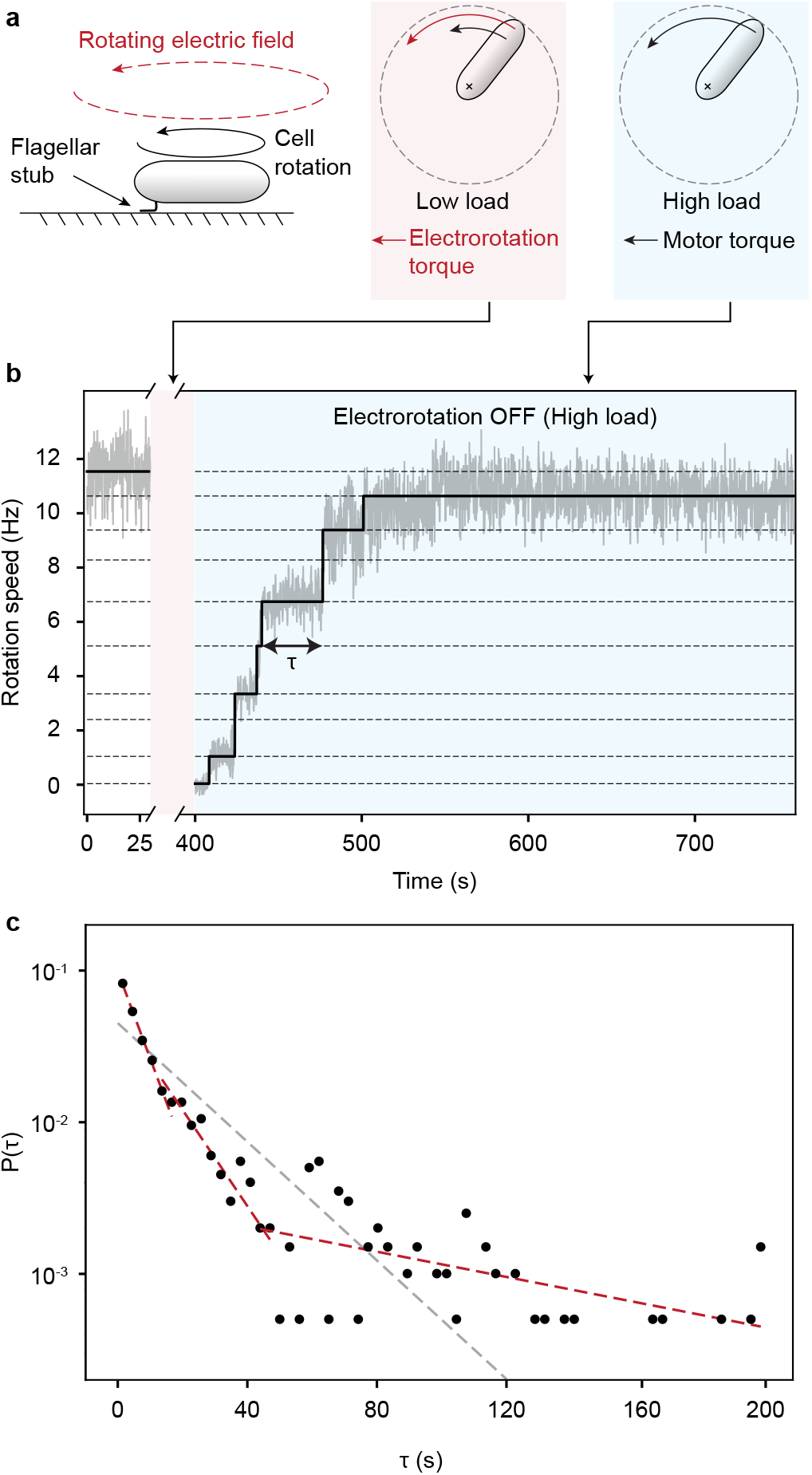
**a**. Experimental strategy. The cell is tethered to a surface via a short flagellar stub (left). A high-frequency rotating electric field then applies an external torque (red) on the cell, which decreases the motor load (middle). Under low load, the motor releases its bound stator units. The electrorotation torque is then switched OFF, which increases the load on the motor (right). In response, the motor recruits stator units, leading to stepwise increases in rotation speed. **b**. Motor speed (gray) as a function of time in an electrorotation experiment, showing fitted steps (black). The dashed lines indicate the discrete speed levels. **c**. Distribution of the dwell times *τ* from the electrorotation experiments (bin size *dτ*=3 s). The dashed red lines are obtained by fitting the data with three exponential functions. The dashed gray line is obtained by fitting the data with a single exponential function.

The time trace of the motor output (i.e., rotation speed) consists of discrete levels that correspond to a given number of stator units driving the motor (**Fig. 1b**). We used a step-fitting algorithm (see *Methods and Materials*) to determine these discrete levels from the noisy speed data at each time instant (black line in **Fig. 1b**). Once the levels were identified, we associated them with a corresponding number of stator units, assuming that when the speed is close to 0 Hz no units are bound to the rotor. Each new higher level is then due to the addition of a single unit. To validate our analysis pipeline, we plotted the entire set of extracted speed levels against the number of stator units associated with those levels (**Fig. S1a**). The approximately linear dependence of rotation speed on the number of bound stator units is consistent with the generally accepted view that, under high load, each stator unit supplies the same torque, resulting in uniform spacing between successive speed levels^14,26^. In addition, we plotted the distribution of the maximum number of stator units (*N*_max_) observed during the remodeling process (**Fig. S1b**). In agreement with previous observations in *E. coli*, we find that the number of stator units bound to the motor is at most 11^27^. The variation in *N*_max_ could be caused by insufficient observation time or different packing arrangements of stators around the circumference of the rotor.

The simplest possible model for the molecular interactions underlying stator remodeling is one in which a stator unit can exist in one of only two states - an unbound state, in which the stator unit diffuses freely in the inner membrane, and a bound state, in which it interacts with the rotor to generate torque. Indeed, this simple model has successfully described the population-averaged kinetics of stator remodeling and its dependence on experimental parameters, such as load^20–22^. This two-state model makes a specific prediction for the dynamics of stator binding and unbinding events that, to our knowledge, has not been tested. It predicts that the dwell time of the stator complex at any given level will be exponentially distributed (see SI). To test this prediction, we plotted the distribution of dwell times at all stator unit numbers (**Fig. 1c**). Evidently, a single exponential function (and therefore a two-state model) is incompatible with the experimental data. The distribution contains at least three exponentially decaying functions. Therefore, a complete model of stator dynamics must contain additional states.

### A mechanistic model for stator dynamics

Motivated by the multiple timescales observed in the distribution of dwell times, we propose a minimal model for stator binding that provides a mechanistic understanding of stator dynamics. Our model is based on the assumption that the affinity between the rotor and a single stator unit is not constant. As a simple approximation of this assumption, we define two separate states for each bound stator unit, characterized by different rates of unbinding. Addition of these internal states explains the emergence of additional time scales in the distribution of dwell times, in line with the experimental evidence.

**Fig. 2a** illustrates the key elements of our model. Away from the motor, a stator unit is in the diffusive state (**D**) in which it floats freely in the inner membrane and the proton channels inside it are closed. When a diffusing stator unit collides with a rotor, it may transition to a bound state (**Fig. 2a**), in which it is tethered to the peptidoglycan (PG) layer of the cell wall and the proton channels open. Subsequently, protons flow through the stator unit and generate torque (Γ) that drives the rotation of the rotor, which has a radius *R*. Newton’s third law dictates that a counter force *F* = Γ*/R* acts on the stator unit in the opposite direction (**Fig. 2a**). This counter force moves the tethered stator unit away from its landing point by a displacement that depends on the force *F*, and thus on the torque Γ. We hypothesize that the rate at which the stator unit unbinds from the rotor decreases with increasing displacement from the landing point (see SI for details). To simplify the problem, instead of modeling a continuous displacement that depends on the torque, we use a coarse-grained approximation (**Fig. 2b**) to describe the internal state of a bound stator unit as being in one of two possible states: a loosely bound (**L**) state with a higher unbinding (off) rate *k*_off,l_, and a tightly bound (**T**) state with a lower off rate *k*_off,t_ ≪ *k*_off,l_. The loose state (**L**) corresponds to a stator unit with a small displacement; the tight state (**T**) represents a stator unit with a large displacement.

**Figure 2.**
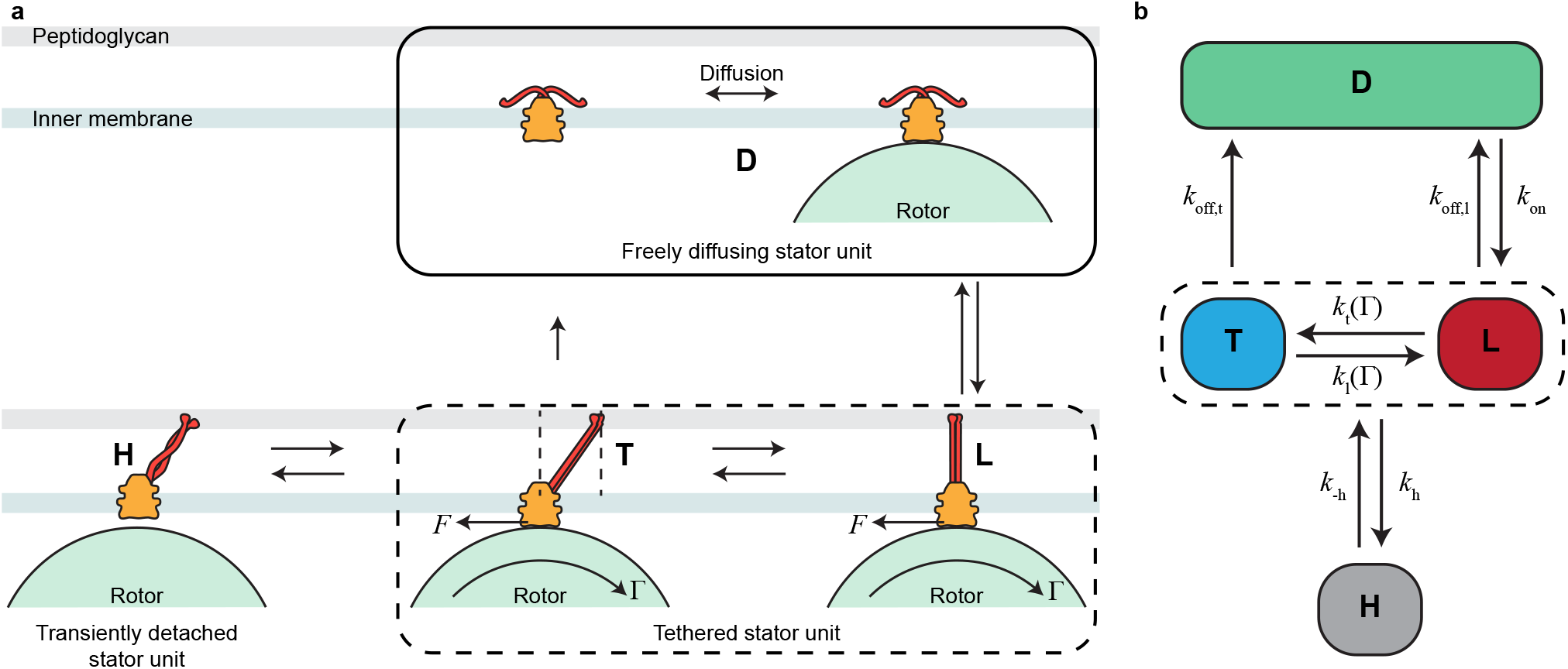
Illustration of the model. **a**. An unbound stator unit is in a closed state (**D**) in which it freely diffuses within the inner membrane and does not translocate protons. Upon interacting with a rotor of radius *R*, the stator unit transitions into a loosely bound state (**L**). In this state, the protons start flowing through the stator unit, resulting in the production of a torque Γ on the rotor. As a reaction, a force *F* = Γ*/R* acts on the stator unit, which displaces it in the direction of the force. This displacement away from the point of tether lowers the off rate and results in a tightly bound state (**T**). A bound stator unit can occasionally detach from the rotor, resulting in a transient ‘hidden’ state (**H**). **b**. A simplified 4-state model of stator dynamics. A stator unit can go between the diffusive state (**D**) and the loosely bound state (**L**) with an on rate, *k*_on_ and an off rate *k*_off,l_. From the **L** state, it can go to the tightly bound state (**T**) with a rate *k*_t_ and back with a rate *k*_l_, both of which depend on the torque (Γ). From the **T** state, it can transition to the **D** state at a much lower off rate *k*_off,t_. This model also includes the transiently detached hidden state (**H**). The **H** state couples with either the **T** or the **L** state with rates *k*_h_ and *k*_−h_.

This coarse-grained model has two important features that dictate its behavior. The first is that, when a freely diffusing stator unit in the **D**-state binds the motor, it enters the loose state (**L**) by definition, at a rate *k*_on_. The on rate for the entire complex, *k*_+_, depends upon the number of available sites (*N*_tot_ − *N*), where *N*_tot_ is the total number of sites and *N* is the number of occupied sites. It can additionally depend on other parameters, such as the motor speed *ω*, which is linearly proportional to the number of attached stator units *N* (**Fig. S1a**). The second important feature of this model is that the transition rates (*k*_l_ and *k*_t_) between the loose state (**L**) and the tight state (**T**) depend on the torque (Γ). In particular, 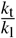 increases with torque so that the equilibrium probability of a bound stator unit to be in the tight state *P*_t_ = *k*_t_*/*(*k*_t_ + *k*_l_) increases with torque. Consequently, the effective (observed) off rate 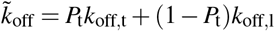, which is averaged over the two bound states (**T** and **L**), is smaller at higher torque because the tight state has a much smaller off rate *k*_off,t_(≪ *k*_off,l_), which is set to zero (*k*_off,t_ = 0) hereafter for simplicity. Thus, torque controls the effective off rate by controlling the internal dynamics of the bound stator units.

While the above described states (**D, L**, and **T**) may be sufficient to describe the on process from freely diffusion unbound stator units and the torque dependence of the off process, our data as well as previous study^25^ suggest that there is an additional “hidden” unbound state different from **D**. Specifically, a bound stator unit can occasionally become detached from the rotor at a rate *k*_h_. The detached unit quickly attaches again to the rotor with a much higher rate *k*_−h_ ≫ *k*_h_, resulting in this hidden state being very short lived. To account for this finding, we introduce this hidden state (**H**) into our model (**Fig. 2b**). The **H**-state is an unbound state (it does not exert force on the rotor). However, it is also not the same as the diffusive state (**D**) because the stator unit stays in the vicinity of the rotor and quickly rebinds with the rotor and becomes attached again. The short-lived **H** states are evidenced by the fastest decay time scale in the distribution of dwell times (the first dotted red line in **Fig. 1c**).

Altogether, we have the minimal 4-state (**D-L-T-H**) model that describes the dynamics of stator assembly (**Fig. 2b**). Aside from the rates (*k*_±*h*_) related to the fast transient state (**H**), the model is defined by four rate parameters: *k*_on_, *k*_off,l_, *k*_t_, and *k*_l_. The on rate *k*_on_ can additionally depend on *N* because of its dependence on the rotation speed *ω*, which is linearly proportional to *N* (**Fig. S1a**). These important biophysical parameters are hard to measure directly. In the following, we determine their values from a statistical analysis of the remodeling data obtained from single flagellar motors during their adaptation to a sudden increase in load.

### The statistics of dwell times and first-passage-time analysis

Due to a separation of timescales, i.e., the transition rate *k*_−h_ from the **H** state to the bound state being faster than the other rates, we simplify our analysis by first identifying the **H** states in the time series using a short time-scale threshold for the duration of the **H** states (see **Fig. S2** in SI for details). From the **H** states identified in the experimental data, the kinetic rates (*k*_±*h*_) to and from the **H** state can be determined.

Once the short-lived **H** states are identified, we can separate them from the rest of the time series and focus on analysing the transitions between the other three states (**L, T**, and **D**), which enable mechano-adaptation in the flagellar motor. An *N*-stator state ends with either an increase or decrease in *N* (*N* → *N* ± 1). The stochastic dynamics of *N* are controlled by both the on (‘+’) and off (‘-’) processes, and whether the ‘+’ or the ‘-’ transition is observed depends on which of the two happens first. Thus, the statistics of dwell times can be understood as a first-passage-time (FPT) problem^28^ with two independent stochastic processes (‘+’ and ‘-’). Mathematically, when the number of stator units reaches *N* at time *t* = 0, we can compute the survival probability *S*(*t*), i.e., the probability the motor stays in state-*N* at a later time *t* ≥ 0. The distribution *P*(*τ*) for dwell time *τ* can then be determined from 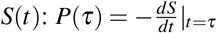

In our model, the on process has a time-independent rate *k*_+_ = (*N*_tot_ − *N*)*k*_on_(*N*), where *k*_on_(*N*) is the on rate for a single stator unit. However, the off process has a rate *k*_−_(*t*|*N*) that is time-dependent because of the existence of the two bound states and their different off rates. This can be understood by considering a unit-*i* (*i* = 1, 2,.., *N*) that binds to the rotor at time *t*_*i*_(≤ 0) in the **L** state. The survival probability *S*_i_(*t*) can be determined analytically:

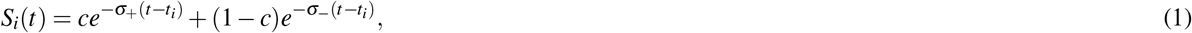

where 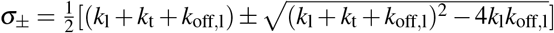 are the two eigenvalues of the transition rate matrix for the two bound states (**T** and **L**), and 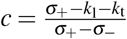 is a constant (see SI for detailed derivation). The off rate for the unit-*i* can then be determined from *S*_*i*_ for *t* ≥ *t*_*i*_:

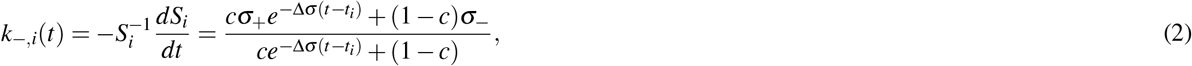

where Δ*σ* = *σ*_+_ − *σ*_−_. Equation 2 shows that *k*_−,*i*_ starts with the “bare” off rate *k*_−,*i*_(*t*_*i*_) = *cσ*_+_ + (1 − *c*)*σ*_−_ = *k*_off,l_ when the stator unit first becomes bound at *t* = *t*_*i*_ and decreases to its equilibrium value *k*_−,*i*_(∞) = *σ*_−_ at *t* − *t*_*i*_≫ *τ*_*e*_ with *τ*_*e*_ = Δ*σ* ^−1^ is the timescale to reach equilibrium between the two bound states (**T** and **L**).

The overall off rate (summed over all the bound stators), 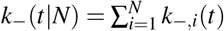, thus also depends on time, which indicates a non-Markovian (memory) effect due to the existence of the hidden bound states ^1^. Given the rates *k*_+_(*N*) and *k*_−_(*t*|*N*), the survival probability satisfies:

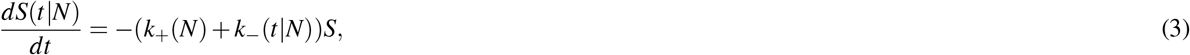

which can be solved with the initial condition *S*(0|*N*) = 1 to obtain a closed-form expression for the survival probability: 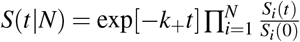. Details of the FPT analysis and derivation of the solution for *S*(*t*|*N*) are given in the SI.

### Quantitative comparison between experiments and theory

First, we estimated the transition rates to and from the **H** state as *k*_h_ ≈ *N*_H_*/T*_tot_ and *k*_−h_ ≈ 1*/τ*_H_, where *N*_H_ is the number of occurrences of the short-lived **H** state, *τ*_H_ is their average dwell time, and *T*_tot_ = Σ *N*_i_*τ*_i_ is a weighted sum of dwell times where each weight is the number of stator units that are already bound. From our experiments, we have *N*_H_ = 43, ⟨*T*_tot_⟩ = 136620 *s*, and *τ*_H_ = 4.1 *s*, leading to *k*_h_ ≈ 0.0003 *s*^−1^ and *k*_−h_ ≈ 0.24 *s*^−1^, which are in excellent quantitative agreement with the rates determined previously by measurements of steady-state motor dynamics^25^.

Next, for each *N*, we separated the statistics for dwell times into two distributions: *P*_+_(*τ*|*N*) and *P*_−_(*τ*|*N*), which are the probabilities for observing the *N* → *N* + 1 and *N* → *N* − 1 transitions, respectively, for the first time at *t* = *τ*, where *t* = 0 is the time when the *N*-state is first reached. The overall distribution of *τ* regardless of the end state is *P*(*τ*|*N*) = *P*_+_(*τ*|*N*) + *P*_−_(*τ*|*N*). The observed histograms for these dwell times (**Fig. S3**) show the occurrence of long dwell times far outside of the exponential distribution with a shorter mean dwell time, which indicates the existence of multiple timescales even for a given *N*. In our model, from the survival probability *S*(*t*|*N*) obtained by solving Eq. 3 for each *N* ∈ [0, *N*_tot_], we can determine the two dwell-time distribution functions *P*_+_(*τ*|*N*) = *k*_+_(*N*)*S*(*τ*|*N*) and *P*_−_(*τ*|*N*) = *k*_−_(*τ*|*N*)*S*(*τ*|*N*), which can be compared with the observed distribution of dwell times.

Because of the limited number of experimental measurements, we compare model and experiments based on lower order moments of the dwell time distributions rather than the full distributions. Specifically, we compute the following statistical properties of the distribution of dwell times for each *N*:

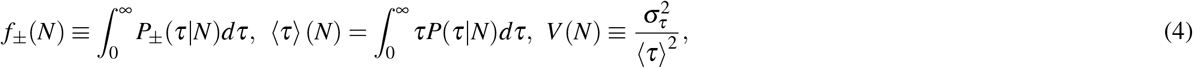

where *f*_±_ is the fraction of ‘+’ (on) or ‘-’ (off) transitions (*f*_+_ + *f*_−_ = 1); (*τ*)(*N*) is the mean dwell time for the *N*-stator state; *V* is the variance of dwell time 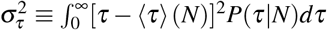 normalized by the square of its mean, which measures the magnitude of the variation in dwell times. We can also define (*τ*_±_)(*N*) = *P*_±_(*τ*|*N*)*dτ* as the average ‘+’ or ‘-’ dwell time, and we have: ⟨*τ*⟩ (*N*) = ⟨*τ*_+_⟩ (*N*) *f*_+_(*N*) + ⟨*τ*_−_⟩ (*N*) *f*_−_(*N*).

In **Fig. 3**, we show the comparison between experimental data and our model results for the average dwell time ⟨*τ*⟩, the fraction of the ‘+’ events *f*_+_, and the normalized variance *V* for different numbers of stator units *N*. The agreement between the experimental data and our model results supports our theory, which incorporates multiple hidden bound states. This is seen most clearly in the normalized variance *V* (**Fig. 3c**). If there were only one bound state, the dwell time distribution would be a single exponential distribution, which would lead to *V* = 1 (see SI). From the experimental measurements, we found ⟨*V*⟩ = 2.0 ± 0.4. The large normalized variance can only be explained by a model with multiple bound states like the one we proposed in **Fig. 2**, which leads to a theoretical value ⟨*V*⟩ = 1.75 ± 0.4, which is consistent with experiments ^2^.

**Figure 3.**
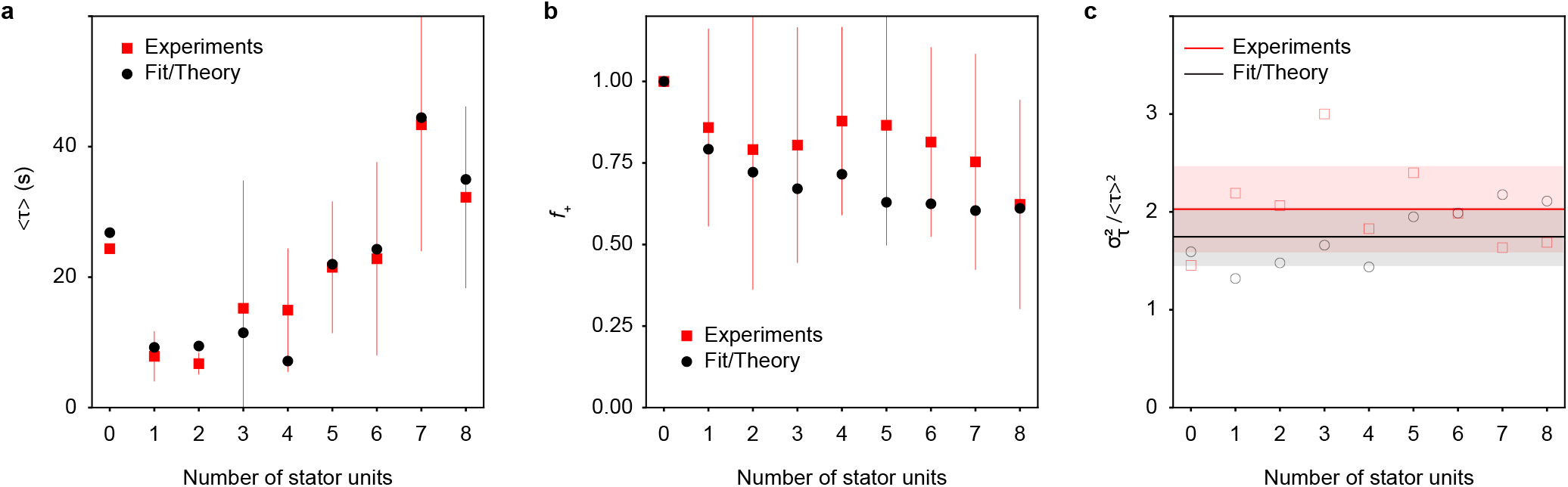
Quantitative comparison between experimental data (red) and theory (black). **a**. The average dwell time ⟨*τ*⟩ for each stator number *N*. **b**. The fraction of on events *f*_+_ for each stator unit number *N*. **c**. The normalized variance 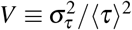 versus *N*. The average *V* over *N* and the standard deviation of *V* over *N* are shown as the solid line and the shaded band for experiments and theory, respectively.

The quantitative agreement between our model and the experimental data confirms the existence of multiple bound states. Furthermore, it allows us to determine the dominant off rate *k*_off,l_ from the **L**-state and the transition rates (*k*_t_, *k*_l_) between the two bound states (**T** and **L**), which are not possible to measure directly. In particular, from the fitting of our model to the experimental data (**Fig. 3**), we determined the values of key model parameters: *c* = 0.30 ± 0.11, *σ*_+_ = 0.19 ± 0.1s^−1^, and an upper bound for the much lower rate *σ*_−_ ≤ 0.0005s^−1^ (see SI for details). In the regime *σ*_+_ ≫*σ*_−_, which is valid at high load, these model parameters are related to *k*_t_,*k*_1_, and 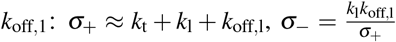, and 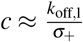, from which we obtain: *k*_off,l_ ≈ *cσ*_+_ = 0.057 ± 0.023*s*^−1^, *k*_t_ ≈ (1 − *c*)*σ*_+_ = 0.13 ± 0.07*s*^−1^, and *k*_l_ ≈ *σ*_−_*/c* ≤ 0.0017*s*^−1^. Thus, the results of our fit show that, in our experiments at a high load, *k*_t_ ≫ *k*_l_, which leads to a much decreased effective off rate *σ*_−_ ≈ (1 + *k*_t_*/k*_l_ + *k*_off,l_*/k*_*l*_)^−1^*k*_off,l_ ≪ *k*_off,l_. The reason for this significant decrease in the off rate at high load is that, once bound in the **L**-state, a stator unit quickly transitions to the more stable **T**-state. The range of equilibrium off rate *σ*_−_ at high load inferred from our experiments is consistent with those measured in previous experiments^25^. At low load, the equilibrium is shifted towards the **L**-state, i.e., *k*_t_ ≪*k*_l_, and our model predicts a much larger effective off rate *σ*_−_ ≈ *k*_off,l_ = 0.057*s*^−1^, which is in excellent agreement with previous measurements at low load^20^.

Our analysis also provides new information about the on process. Based on the FPT analysis, we have *P*_+_(*τ*|*N*) = *k*_+_(*N*)*S*(*t*|*N*), which leads to 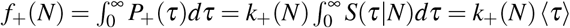. Therefore, we can obtain the on rates for different stator number *N, k*_+_(*N*) = *f*_+_(*N*)*/* ⟨*τ*⟩ (*N*), from the observed *f*_+_(*N*) and ⟨*τ*⟩ (*N*). Furthermore, we can obtain *k*_on_ = *k*_+_*/*(*N*_tot_ − *N*) as the on rate for each empty binding site. In **Fig. 4**, we plot *k*_on_(*N*) for different values of *N* = 0, 1, …, 8. This plot reveals several interesting features about the on process. First, the nucleation rate, or the initial on rate for *N* = 0: *k*_on_(0) ≈ 0.0037 s^−1^ is much smaller than the on rates for *N* ≥ 1. Second, the on rates for *N* ≥ 3 are roughly the same, with *k*_on_(*N* ≥ 3) ≈ 0.0067 s^−1^. Third, the on rates for *N* = 1, 2 are higher than the rates for *N* ≥ 3. The first two features were also reported in a recent study by Ito *et al*^29^. At high load, the motor speed is linearly proportional to *N* and the low initial on rate has been attributed to a possible dependence of *k*_on_ on the rotational speed *ω* in a nonlinear, sigmoid-like fashion^29^. However, as far as we know, the enhanced on rates for *N* = 1 and *N* = 2 have not been reported before. These high rates are related to the short average dwell times for *N* = 1 and *N* = 2 (**Fig. 3a**).

**Figure 4.**
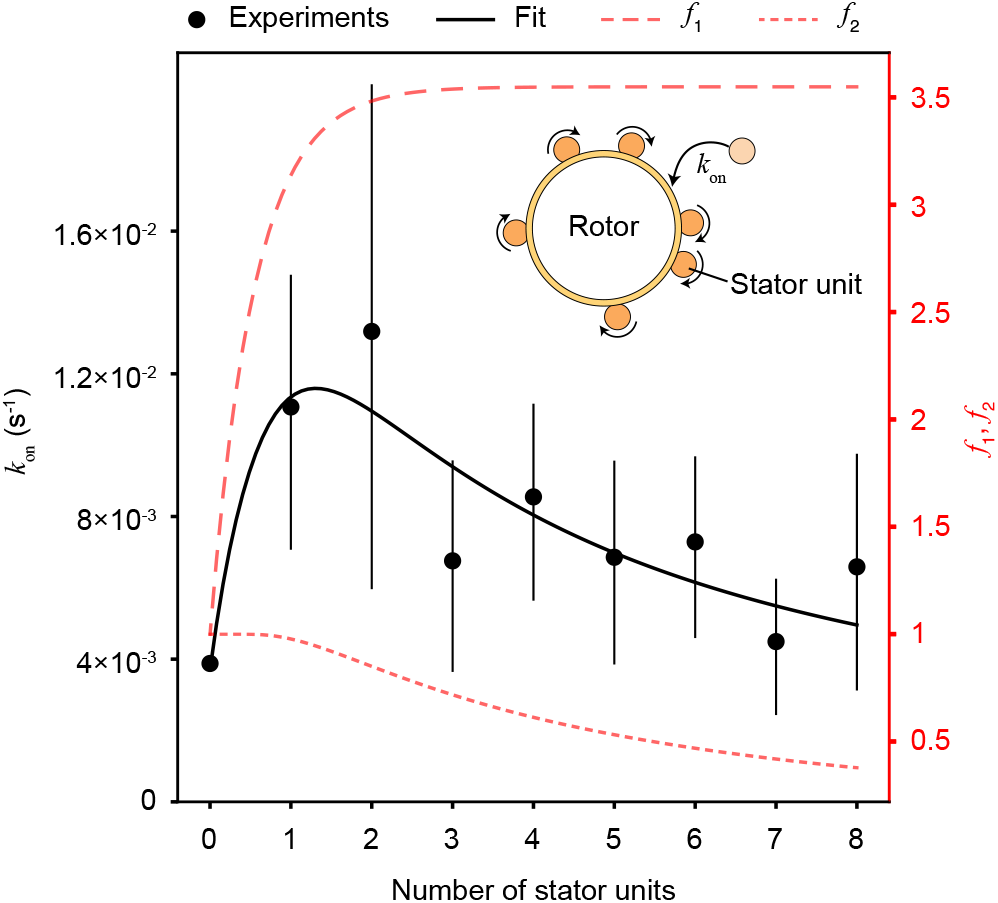
The binding rate *k*_on_ for an individual stator unit at different numbers of bound stator units (*N*). The black curve was obtained from the simple model *k*_on_(*N*) = *k*_on_(0) *f*_1_(*N*) *f*_2_(*N*), where *k*_on_(0) = 0.0037 *s*^−1^ is the on rate for *N* = 0 and the two functions, *f*_1_ and *f*_2_, are given by *f*_1_(*N*) = (1 + *A*(1 exp[*BN*])) and *f*_2_(*N*) = 1 exp[*C/N*]. The parameters *A, B*, and *C* are determined from fitting the experimental data: *A* = 0.23, *B* = 1.8, *C* = 3.8.

We do not understand the molecular details of how a freely diffusing (unbound) stator is incorporated into a rotating motor. The dependence of *k*_on_(*N*) on *N* is likely caused by effect(s) of the motor speed on the on rate. The non-monotonic dependence we found further suggests that there may be multiple counteracting effects. On the one hand, a higher motor speed may enhance both the probability of collision between a diffusive unbound stator unit and the rotor and the collision strength, effects that could contribute to a higher on rate^29^. On the other hand, a higher speed would also mean a shorter contact time, which could lead to a lower on rate, as previously suggested by Wadhwa *et al*^21^. By combining these two possible counteracting effects, we propose a minimal phenomenological model to fit the data:

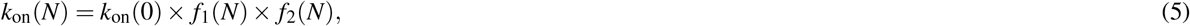

where *f*_1_(*N*) (*f*_2_(*N*)) is a monotonically increasing (or decreasing) function of *N* that describes the speed-dependent enhancing (reducing) effect of the on rate with *f*_1_(0) = *f*_2_(0) = 1. Here, we chose the simplest forms with the smallest number of parameters for these two functions that are consistent with previous studies^21,29^. As shown in **Fig. 4**, this simple model yields reasonable agreement with the experimental results, which support the hypothesis that the motor speed may affect the on process through two opposing mechanisms.

### Summary and Discussion

In this study, we combined quantitative measurements of the remodeling dynamics of single motors and first-passage-time analysis of the statistics of dwell times. We also developed and tested a minimal four-state model that quantitatively captures multiple experimental observations of the remodeling dynamics of the motor. A short-lived unbound state (**H**), proposed in a previous study^25^ of the steady-state motor dynamics, is confirmed in our analysis of the transient dynamics of motor remodeling. We include this **H**-state in our analysis for the sake of completeness, but it is not essential for mechano-adaptation. More importantly, our study reveals the existence of multiple states for a stator unit bound to the motor, each with a different unbinding rate. This multiplicity of bound states is what confers mechano-adaptation to the motor.

In all sensory adaptation systems, a key common feature is the existence of multiple internal states that can be modulated in response to changes in external stimuli^30^. For example, bacterial chemoreceptors have multiple methylation sites that allow chemoreceptors to adapt to changes in ligand concentrations by adjusting their methylation levels^31,32^. The multiple bound states of the stator unit identified here serve as the internal adjustable states necessary for adaptation of the bacterial flagellar motor to changes in external mechanical signals. The transitions between these bound states introduce additional timescales in the transient dynamics of motor remodeling, which allows us both to explain the experimental results and determine the transition rates between them.

Another recent study proposed a model in which the bound state is split into two states with different unbinding rates^33^. Based on population-averaged kinetics of stator remodeling, those authors showed that a three-state model (also termed a ‘two-state catch-bond model’^34^) can explain the asymmetry observed in the timescales of relaxation to steady state from either a large or small number of stator units. However, the authors were unable to distinguish clearly between a three-state (loosely bound, tightly bound, unbound) model and a two-state model (bound, unbound) with a speed-dependent on rate, as previously proposed by Wadhwa *et al*^21^.

Here, by analyzing the statistics of the dwell times of single motors, we conclusively show that a two-state model is incompatible with the experimental observations. We go a step further by including the short-lived hidden state in our model and confirming its presence in our experimental data. These advances are primarily enabled by the detailed analysis of single binding and unbinding events using first-passage-time methods, rather than by looking at population averages. The convergence between our study and that of Perez-Carrasco *et al*.^33^, which investigated different aspects of mechanosensitive stator remodeling (dwell-time statistics vs. relaxation-time asymmetry), makes the model with multiple bound states a strong candidate for future theoretical and experimental work in this field.

The emerging picture for mechano-adaptive remodeling of the bacterial flagellar motor is that the torque generated by bound stator units controls their off rate by modulating the transition rates between the **T**-state (with a low or zero off rate) and the **L**-state (with a high off rate *k*_off,l_), and thus the equilibrium between the two bound states. The ratio 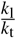 therefore depends on the torque Γ and determines the observed off rate. Quantitatively, we find that the motor can tune its effective off rate over a wide range (*>* 100 fold) from a very low rate (≤ 0.0005*s*^−1^) at high torque (near stall) to a much higher rate (∼ 0.057*s*^−1^) at low torque. Thus, the motor can adapt to changes in the mechanical environment by adjusting the number of stator units over a range of *N*_tot_ from 0 to 11. In our experiments, the motor drove the rotation of tethered cells and therefore operated close to stall. As a result, our data can only be used to determine the values of *k*_*t*_ and *k*_*l*_ at high torque (near stall). In the future, it will be interesting to use the same modeling and analysis framework developed here to analyze the data from experiments performed at different loads - for example, by using flagella labeled with beads of different sizes - to dissect the dependence of *k*_l_ and *k*_t_ on torque.

As first pointed out by Nord *et al*.^22^, the large reduction in the unbinding (off) rate of stator units as torque increases is reminiscent of the general catch-bond phenomenon, in which the lifetime of a receptor-ligand bond increases with tensile force applied to the bond^35^. In the case of stator units, however, the molecular mechanism causing the differences in off rates for different bound states remains unclear. In **Fig. 2**, we suggest a scenario in which the off rate may depend on the physical location (relative to its tethering position) of a bound stator unit, which is affected by the torque it generates. Another possibility is that the binding strength is increased by the torque through mechanically induced allostery in which torque production (and the resultant force) may propagate through the stator unit and cause conformational changes in the peptidoglycan binding domain of the stator unit. Indeed, these are only two of the several possible scenarios that can lead to the catch-bond phenomenon^35^. Currently, there is no evidence to rule out either of these scenarios, both of which can be described by our model with two bound states (**T** and **L**). However, the presence of two bound states makes the binding of the stator unit to peptidoglycan analogous to the catch-bond behavior of FimH-mannose^36,37^, kinetochore-microtubule^38^, and vinculin-F-actin interactions^39^, all of which also consist of two bound states. These contrast with the selectin-ligand^40^ and myosin-actin^41^ interactions, which exhibit a single time scale in their dwell time distributions.

Our study also reveals several interesting features of the dependence of the on rate on the number of stator units. Previously, Wadhwa *et al*. suggested a model in which the on rate decreases with motor rotation speed, which would decrease on rate with an increase in the number of stator units^21^. In contrast, recent work by Ito *et al*. found that the on rate increases with motor rotation speed, specifically when going from *ω*= 0 to *ω* > 0^29^. Our data reveal that the actual behavior is a combination of the two. The observed on rate has a non-monotonic dependence on the number of stator units: the on rate for *N* = 0 is smaller than that for *N* = 1 or 2, where it peaks. At *N* > 2, the on rate decreases. Indeed, a phenomenological model that combines features of these two competing effects successfully captures the observed trend. The non-monotonic dependence of the on rate on the number of bound stator units suggests that the on process may contain two (or more) steps with opposing dependence on the motor rotation speed. In principle, the on rate could depend on the number of bound stator units (via a possible cooperative binding effect) as well as on the motor speed *ω*. However, in our current experiments at high load, *ω* is linearly proportional to *N*, which makes it impossible to separate the dependence on *N* and *ω*. Future experiments that systematically measure the on rate under different loads are needed to understand the on process and its dependence on *N* and *ω*.

A model with multiple bound states with mechanically regulated transition rates, such as the one proposed here for the bacterial stator units, also provides a possible mechanism for downstream signaling of mechanical signals. It is conceivable that, apart from regulating the binding strength of proteins, force could also alter their biochemical interactions with downstream signaling molecules. This could explain the putative role of the bacterial flagellar motor, and stator units in particular, in surface sensing during biofilm formation and differentiation of swarmer cells^42–45^.

Overall, this work demonstrates the power of combining quantitative data from single molecule experiments with detailed stochastic analysis (e.g. first-passage-time analysis) to decipher the underlying mechanisms in biological systems without requiring that all the molecular details be known. Similar approaches should be applicable to studying other stochastic dynamic processes in biology, especially those that have to do with self-assembly of multi-protein complexes and their regulation by intrinsic or extrinsic signals.

## Methods and materials

### Bacterial strains and cultures

We used *Escherichia coli* strain KAF95 (alias HCB986; a derivative of AW405) for all experiments. This strain is deleted for the chemotaxis response regulator CheY. Consequently, the cells of this strain rotate their flagellar motors exclusively counterclockwise. Additionally, this strain is deleted for the WT flagellin gene FliC and transformed with the plasmid pFD313, which expresses sticky FliC, resulting in filaments that readily attach to a variety of surfaces. Cells grown at 33 °C to OD_600_ =0.5 - 0.6 were washed and resuspended in TES buffer (20 mM TES, 0.1 mM EDTA, pH = 7.5). The cells were then sheared by passing through a 20 cm long piece of polyethylene tubing (inner diameter 0.50 mm) 60 times. The cells were then washed again to remove the sheared filaments and re-suspended in TES buffer.

### Electrorotation experiments

The electrorotation apparatus has been described before^21,46^. Cells were introduced into a custom-built flow cell that consisted of a circular sapphire window on one side and a circular glass cover slip on the other side. Cells readily tethered to sapphire via a short flagellar stub, which resulted in rotation of the cell body around the point of tether. The flow cell also contained four tungsten micro-electrodes whose tips were located at a short distance from the sapphire surface. The electrodes were driven in quadrature at 2.25 MHz to apply a rotating electric field on the tethered cells. This field caused an external torque on the cell body in the same direction as the torque applied by the flagellar motor. The strength of the external torque could be tuned by changing the amplitude of the rotating electric field. The temperature of the sapphire window was held constant at 20 °C by a circular Peltier element driven by a proportional controller. The flow cell and electrode assembly was fixed on the 20X objective of a phase contrast microscope. Rotating cells were imaged at 50 or 100 frames per second using a high-speed sCMOS camera (Edge 5.5; PCO-Tech).

### Data analysis and step fitting

We measured the angular displacement of the cell body between consecutive frames and multiplied it with the imaging frame rate, followed by filtering with a median filter of order 15. This provided the rotation speed of the motor as a function of time (gray line in **Fig. 1**). We then proceeded to fit steps to the rotation speed to extract the number of active stator units as a function of time. Of all the traces, we selected only those for further analysis in which the rotation speed was below 1 Hz at the end of the electrorotation phase, so that at most a single stator unit was bound to the motor at the beginning of the high load phase.

Our approach to step-fitting consists of partitioning the total time in smaller time intervals in such a way that we minimize the point-by-point distance between the fitted step and the original trace. We call *ω*(*t*_*i*_) the value of the rotation speed at time instant *t*_*i*_ from the original data and, given that the time increment *δt* is constant for every step, we have *t*_*i*_ = *i* * *δt*. We will call *t*_*n*_ the total time of a given experiment, where *n* is the total number of measurements per trace.

We need to find an instant *t*_*l*_ at which a step occurs. The point of the partition (the index *l*) is chosen in such a way that the updated step function will be as close as possible to the original data. More formally, we define the averages

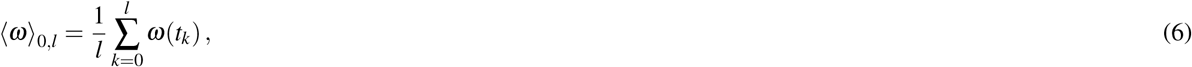

and

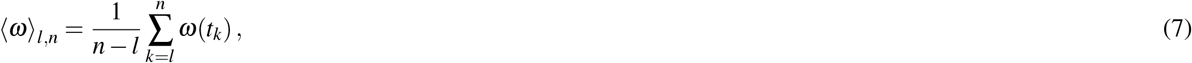

where ⟨*ω*⟩_0,*l*_ is the average of the original data in the time interval between *t*_0_ = 0 and *t*_*l*_ and ⟨*ω*⟩_*l,n*_ is the average of the data in the interval between *t*_*l*_ and the final instant *t*_*n*_. Then, *l* is chosen in such a way that the residual Δ, defined as sum of the squares of the difference between the fitted step and the original data at each time instant:

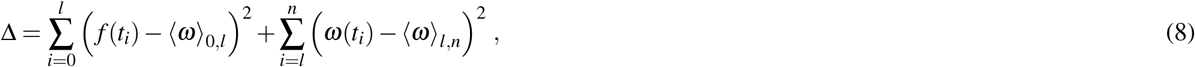

is minimal. The partition is then iterated several times. The formula for Δ after *K* iterations is

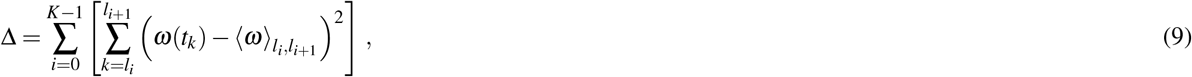

where *l*_*i*_ is the index for the instant of at which the *i*-th step in the fitted function occurs and the indices are defined so that *l*_*i*_ < l_*i*+1_ for all values of *i* (*i* = 0, 1, …, *K* − 1).

The choice of the total number of iterations *K* is important in order to avoid over-fitting. To determine when to interrupt the loop, at each iteration we calculate the average 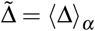 that is obtained by partitioning at randomly generated instants *t*_*α*_ rather than the specific instants that minimize Δ to Δ_min_. Iteration is stopped as soon as 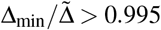, i.e., when compared to an equal number of randomly placed steps, an additional computed step would not improve the fit by more than 0.5%.

The process described above results in a fitted step function *g* for a given time *t*_*u*_, where *l*_*i*_ < *u* < *l*_*i*+1_:

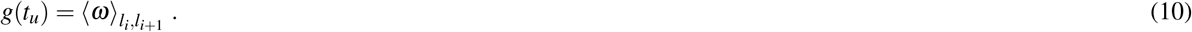

At this stage, we must associate the discrete values of *g* to a specific number of stator units. To do so, we determine whether slightly different values of *g* during different time intervals correspond to the same number of stator units. This is done with a sorting method. First, we sort all the values of *g* in ascending order. Then, if two consecutive values of *g* (levels) differ by less than 0.75 Hz, we assign to them a new level given by the weighted average of their current levels. Then, we associate a number of stator units *N* to each level starting with *N* = 0 for the level with a value close to 0 Hz and we add a stator unit for each subsequent level.

## Acknowledgements

We are grateful to Prof. Mike Manson for his critical reading of the manuscript. This work was supported by the National Institutes of Health under Award Numbers K99GM134124 (to N.W.) and R35GM131734 (to Y.T.).

## Footnotes

1 The T-state and the L-state are “hidden” states because we can not tell them apart directly from the experimental observations, i.e., torque (or speed) of the motor.

2 The variability in *V* (*N*) for different *N* is likely due to the relatively small sample size and the fact that *V* is a higher order measure of the the statistics of dwell times.

