## Supplementary Information for "A multi-state dynamic process confers mechano-adaptation to the bacterial flagellar motor"

**Navish Wadhwa**[1,✉], **Alberto Sassi**[2], **Howard C. Berg**[1], **and Yuhai Tu**[2,✉]

[1]Department of Molecular and Cellular Biology, Harvard University, Cambridge, MA 02138

[2]IBM T. J. Watson Research Center, Yorktown Heights, New York, NY, 10598, USA

### Supplementary Figures

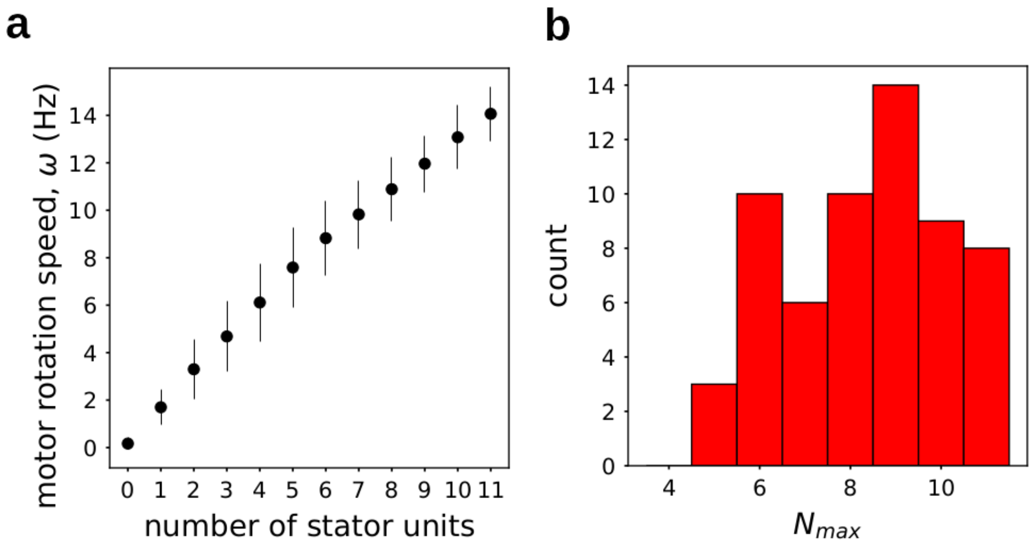


**Figure S1.** **a.** Average rotation speed vs number of stator units. Vertical lines represent standard deviation. **b.** Histogram of the largest number of bound stator units reached in an experiment. We have $N_{\text{max}} \leq N_{\text{tot}}$, where $N_{\text{tot}} = 11$ is the maximal number of stator units that can be accommodated in a flagellar motor.

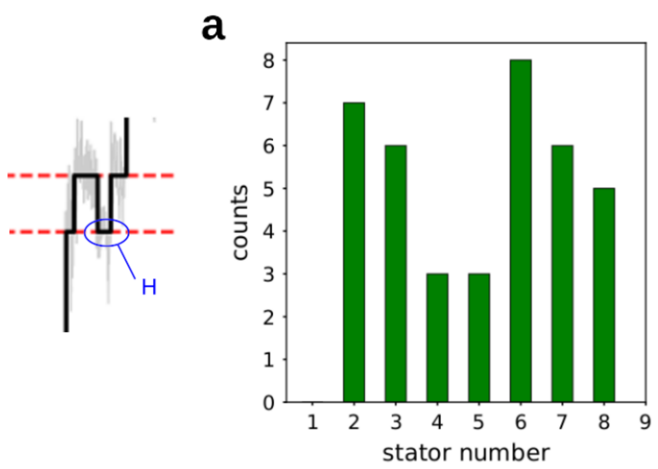


**Figure S2.** Occurrence of short "wells" for different stator numbers due to transition to the short-lived unbound state. A short well is defined as an unbinding event followed by a binding event within $\Delta\tau < 8$ s. **a.** Counts of short wells observed at different stator units numbers. **b.** Total counts of wells observed for different values of the threshold duration ($\delta\tau$) of the well.

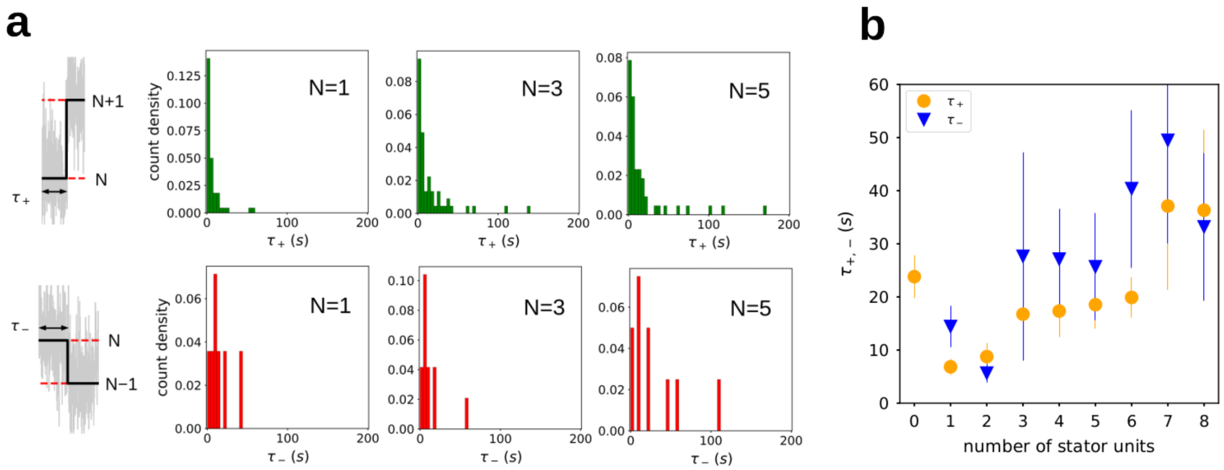


**Figure S3.** **a.** Histograms of dwell times ending with binding (above) and unbinding (below) of a new stator unit, for three different values of $N$. **b.** Conditional dwell times $\tau_+$ and $\tau_-$ from the electrorotation experiments vs the number of stator units bound to the rotor.

### Supplementary text

#### A plausible model for stator dynamics and its simplified form

Here we present a conceptual model to described the stochastic motion of a bound stator unit to demonstrate a plausible mechanism for the torque-dependence of the off rate. The basic idea is that a bound (tethered) stator unit can become unbound with a unbinding rate $k_{\mathrm{off}}(x)$ that can depend on its displacement $x$. This dependence of $k_{\mathrm{off}}$ on $x$ is reasonable as there are barriers between the bound and unbound states that can depend on location of the stator unit relative to the landing point (the attachment point at $x=0$). We assume that $k_{\mathrm{off}}(x)$ is maximum at $x=0$ as the stator unit can unbind through the same pathway as the binding process at $x=0$ and it decreases as $|x|$ increases, e.g., $k_{\mathrm{off}}=k_0\exp(-|x|/x_0)$ or $k_{\mathrm{off}}=k_0\exp\left(-x^2/x_0^2\right)$. This symmetric form of $k_{\mathrm{off}}$ for $x\to -x$ is inspired from the CCW and the CW symmetry in stator remodeling[1].

For a tethered stator unit, we can write down the dynamics of its displacement $x$ as a Langevin equation:

$$\frac{dx}{dt}=\xi^{-1}\left(-\frac{dF_0(x)}{dx}-f\right)+\eta, \tag{S1}$$

where $\xi$ is the viscosity; $\eta$ is the thermal noise with $\langle\eta\rangle=0$ and $\langle\eta(t)\eta(t')\rangle=2k_BT\xi^{-1}\delta(t-t')$ ($k_BT$ is the thermal energy unit); $F_0(x)$ is the free energy of the tethered stator. If we assume the tethers can be approximated as springs with a spring constant $k$, we have: $F_0(x)=\frac{1}{2}kx^2$. Of course, $F_0(x)$ can have other symmetric form such as $F_0(x)=k|x|$ or even have other local minima than $x=0$.

The driving force $f=\Gamma/R$ is due to the counter force from the rotor to the stator unit with $\Gamma$ the torque and $R$ the rotor radius. If we use the long time approximation and consider $\Gamma$ as a constant, We can solve Eq. S1 and obtain the steady state distribution of $x$ as:

$$P_s(x)=A\exp[-(F_0(x)+\Gamma x/R_0)/k_BT], \tag{S2}$$

where $A$ is normalization constant. Given the steady state distribution of $x$ given in Eq. S2 and the dependence of the off rate on $x$ given by $k_{\mathrm{off}}(x)$ to determine the observed overall off rate $\tilde{k}_{\mathrm{off}}$:

$$\tilde{k}_{\mathrm{off}}(\tau)=\int P_s(x)k_{\mathrm{off}}(x)dx, \tag{S3}$$

which depends on $\Gamma$ because $P_s(x)$ depends on $\Gamma$.

For simplicity, we can coarse grain the continuous $x$-dependence by two states, **T** and **L**, where the two states are separated by a displacement threshold $x_{\mathrm{th}}$. **L** represents a loose state with $|x|\le x_{\mathrm{th}}$ around $x\simeq 0$ where $k_{\mathrm{off}}(x)\equiv k_{\mathrm{off,l}}$ is large, and **T** represents a tight state at $|x|>x_{\mathrm{th}}$ where $k_{\mathrm{off}}(x)\equiv k_{\mathrm{off,t}}$ is much smaller than $k_{\mathrm{off,l}}$. This simplified model of the bound stator is used in this work.

#### The duration time statistics by using first-passage time analysis

Assume at $t = 0$, the motor enters into the state-$N$ with $N$ engaged stator units (by either adding or removing a stator unit). The motor exits the state-$N$ by either having a new stator unit engaged with the rotor (the “on" process) or by having one of the $N$ bound stator units leaving the motor (the “off" process), whichever happens first.

Since the observed transition (either on or off) depends on which transition happens first, the problem is a first passage time problem. The first passage time problem can be treated by computing the survival probability $S(t)$, which is the probability the motor stays in state-$N$ at time $t \geq 0$. Obviously, we have $S(0) = 1$ and $S(\infty) = 0$.

There are two processes (on and off) that terminate the state-$N$. The on process has a constant (time-independent) rate:

$$k_{+} = (N_{\mathrm{tot}} - N)k_{\mathrm{on}}, \tag{S4}$$

where $N_{\mathrm{tot}}$ is the maximum number of stator units each motor can accommodate, and the on rate $k_{\mathrm{on}}$ can depend on the rotational speed $\omega$, which is linearly proportional to the number of stator units ($\omega = \omega_1 N$ in the high load regime with $\omega_1$ the rotational speed per stator unit). Additionally, $k_{\mathrm{on}}$ can also depend directly on $N$ due to other mechanisms, such as co-operative interactions.

For $N \geq 1$, the off process has a rate $k_{-}(t)$ that is more complex as there are multiple stator units each with a different effective “off” rate depending on their arrival time $t_N < t_{N-1} < .. < t_1 \leq 0$ where the newest stator unit arrives at $t_1 \leq 0$. For a given unit-$i$, the probability of it being still bound with the rotor is its “survival” probability $S_i(t)$. The survived unit-$i$ can be either in the **L** or the **T** state with probability $L_i(t)$ and $T_i(t)$, respectively, and $S_i = T_i + L_i$. $L_i(t)$ and $T_i(t)$ can be determined by solving the following equations:

$$\frac{dL_i}{dt} = -(k_{\mathrm{off,l}} + k_{\mathrm{t}})L_i + k_{\mathrm{l}}T_i, \tag{S5}$$

$$\frac{dT_i}{dt} = -k_{\mathrm{l}}T_i + k_{\mathrm{t}}L_i, \tag{S6}$$

with the initial conditions $L_i(t_i) = 1$ and $T_i(t_i) = 0$ as we assume the stator is in the L state when it initially binds to the rotor.

The Eqs. S5&S6 can be solved to obtain:

$$L_i(t) = \frac{\sigma_{+} - k_{\mathrm{l}}}{\Delta\sigma} e^{-\sigma_{+}(t-t_i)} + \frac{k_{\mathrm{l}} - \sigma_{-}}{\Delta\sigma} e^{-\sigma_{-}(t-t_i)}, \tag{S7}$$

$$T_i(t) = \frac{k_{\mathrm{t}}}{\Delta\sigma}(e^{-\sigma_{-}(t-t_i)} - e^{-\sigma_{+}(t-t_i)}), \tag{S8}$$

$$S_i(t) = T_i(t) + L_i(t) = \frac{\sigma_{+} - k_{\mathrm{l}} - k_{\mathrm{t}}}{\Delta\sigma} e^{-\sigma_{+}(t-t_i)} + \frac{k_{\mathrm{l}} + k_{\mathrm{t}} - \sigma_{-}}{\Delta\sigma} e^{-\sigma_{-}(t-t_i)}, \tag{S9}$$

where $\sigma_{\pm} = \frac{1}{2}[(k_{\mathrm{l}} + k_{\mathrm{t}} + k_{\mathrm{off,l}}) \pm \sqrt{(k_{\mathrm{l}} + k_{\mathrm{t}} + k_{\mathrm{off,l}})^2 - 4k_{\mathrm{l}}k_{\mathrm{off,l}}}]$ are the two rate constants (eigenvalues of the rate matrix from Eqs. S5&S6) with $\sigma_{+} > \sigma_{-} > 0$ and $\Delta\sigma \equiv \sigma_{+} - \sigma_{-} > 0$.

Since a stator unit only has a non-zero off rate $k_{\mathrm{off,l}}$ when it is in the L state (i.e., $k_{\mathrm{off,t}} = 0$), the overall off rate $k_-$ is:

$$k_-(t) = k_{\mathrm{off,l}} \sum_{i=1}^{N} p_i(t) = -\sum_{i=1}^{N} S_i^{-1} \frac{dS_i}{dt}, \tag{S10}$$

where $p_i(t) = \frac{L_i}{L_i + T_i}$ is the fractional probability of unit-$i$ to be in the L state at time $t$.

Given the rates $k_+$ and $k_-$, the survival probability satisfies: $S(t+dt) = S(t) \times (1 - (k_+ + k_-)dt)$. By taking the limit $dt \to 0$, we have:

$$\frac{dS}{dt} = -(k_+ + k_-(t))S, \tag{S11}$$

which can be solved with the initial condition $S(0) = 1$:

$$S(t|N) = \exp[-k_+ t] \prod_{i=1}^{N} \frac{S_i(t)}{S_i(0)}, \tag{S12}$$

which is the product of the survival probabilities of all the individual bound stator units and the survival probability ($\exp(-k_+ t)$) due to adding a stator unit. Note that in the product we have $\frac{S_i(t)}{S_i(0)}$ to satisfy $S(0) = 1$.

From Eq. S9, the expression for $S_i$ can be written as:

$$S_i(t) = c\exp(-\sigma_+(t - t_i)) + (1 - c)\exp(-\sigma_-(t - t_i)), \tag{S13}$$

where $c = \frac{\sigma_+ - k_{\mathrm{l}} - k_{\mathrm{t}}}{\sigma_+ - \sigma_-}$ is a constant. From the expression of $S_i$, we can obtain the expression for $\frac{S_i(t)}{S_i(0)}$:

$$S_i(t)/S_i(0) = c_i \exp(-\sigma_+ t) + (1 - c_i)\exp(-\sigma_- t) = e^{-\sigma_- t}[(1 - c_i) + c_i e^{-\Delta\sigma t}], \tag{S14}$$

where the coefficient $c_i$ is given by:

$$c_i = \frac{c\exp(\sigma_+ t_i)}{c\exp(\sigma_+ t_i) + (1 - c)\exp(\sigma_- t_i)} = \frac{ce^{\Delta\sigma t_i}}{ce^{\Delta\sigma t_i} + 1 - c}, \tag{S15}$$

where $\Delta\sigma \equiv \sigma_+ - \sigma_- > 0$. It is easy to see that $c_i < 1$ and since $c$ can be negative, $c_i$ can also be negative. Given the order of the stator unit arrival times: $t_N < t_{N-1} < \ldots < t_1 \leq 0$, the absolute value $|c_i|$ decreases with $i$ (in fact, $|c_i|$ decreases exponentially with $|t_i|$).

Finally, the expression for $S(t|N)$ can be written as:

$$S(t|N) = e^{-(k_+ + N\sigma_-)t} \sum_{n=0}^{N} a_n e^{-n\Delta\sigma t}, \tag{S16}$$

which contains $(N+1)$ exponential decay terms each with a different decay rate (or timescale). The coefficients are: $a_0 =$

$\prod_{i=1}^{N}(1-c_i)>0$, $a_1=\sum_{i=1}^{N} c_i\prod_{j\neq i}(1-c_j)$, $a_2=\sum_{i=1}^{N}\sum_{j\neq i} c_i c_j \prod_{k\neq i,k\neq j}(1-c_k)$, etc. Given that $c_i$ can be negative, $a_1$ can be negative. In principle, $a_n$ can depend on $N$.

Note that Eq. S16 with $(N+1)$ timescales is exact. Given the fact that $\Delta\sigma>0$ and $t_i<0$, we have $|c_N|<|c_{N-1}|<...<|c_1|<1$. For simplicity, we can keep only the larger values of $c_i$ with $i\leq i_m$ and let the other values of $c_i$ to be zero $c_{i>i_m}=0$. For example, the simplest model corresponds to $i_m=1$ and we have:

$$S(t|N)=(1-c_1)e^{-k_a t}+c_1 e^{-k_b t},\quad N\geq 1 \tag{S17}$$

where $k_a=k_+ + N\sigma_-$, $k_b=k_a+\Delta\sigma$, and $S(t)=e^{-k_{+,0}t}$ for $N=0$. The model then contains three parameters $\sigma_\pm$ and $c_1\approx c$ if we assume $t_1\approx 0$.

In this paper, we used $i_m=3$, i.e., by making the approximation that $c_{i\geq 4}=0$, which leads to an explicit expression for $S(t|N)$:

$$S(t|N)=\begin{cases}(1-c_1)e^{-k_a t}+c_1 e^{-k_b t}, & N=1\\ (1-c_1)(1-c_2)e^{-k_a t}+(c_1+c_2-2c_1c_2)e^{-k_b t}+c_1c_2e^{-k_c t}, & N=2\\ (1-c_1)(1-c_2)(1-c_3)e^{-k_a t}+[c_1(1-c_2)(1-c_3)+c_2(1-c_1)(1-c_3)+c_3(1-c_1)(1-c_2)]e^{-k_b t}+ & \\ +[c_1c_2(1-c_3)+c_1c_3(1-c_2)+c_2c_3(1-c_1)]e^{-k_c t}+c_1c_2c_3e^{-k_d t}, & N\geq 3\end{cases} \tag{S18}$$

where $k_c=k_a+2\Delta\sigma$ and $k_d=k_a+3\Delta\sigma$. Note that the normalization condition $S(0)=1$ is preserved in the expression for $S(t)$ given above, which have four decay rates (terms) when $N\geq 3$ and there are three variables $c_1$, $c_2$, and $c_3$ that we used to fit the experimental data. Given that the arrival time for the last stator unit $t_1=0$ (at least for most cases), we have: $c_1\approx c$.

From the expression of $S(t|N)$ given in Eq. S18, we can obtain the expressions for $k_-(\tau|N)=-\frac{d\ln S}{dt}|_{t=\tau}-k_+$ and the two dwell time distributions $P_+(\tau|N)$ and $P_-(\tau|N)$ for the '+' and '-' transitions when the bound stator number is $N$:

$$P_+(\tau|N)=k_+(N)S(\tau|N),\quad P_-(\tau|N)=k_-(\tau|N)S(\tau|N)\,. \tag{S19}$$

The overall dwell-time distribution $P(\tau|N)=P_+(\tau|N)+P_-(\tau|N)$. These dwell time distributions can be used to derive expressions for $\langle\tau\rangle$, $f_+$, and $V$ for each $N$:

$$\langle\tau\rangle(N) = \int_0^\infty \tau P(\tau|N)d\tau\,, \tag{S20}$$

$$f_+(N) \equiv \frac{\int_0^\infty P_+(\tau|N)dt}{\int_0^\infty P(\tau|N)d\tau}\,, \tag{S21}$$

$$V(N) \equiv \frac{\int_0^\infty \tau^2 P(\tau|N)d\tau}{(\int_0^\infty \tau P(\tau|N)d\tau)^2} - 1\,, \tag{S22}$$

which depend on $k_+(N)$, $\sigma_-$, $\Delta\sigma$, $c_1$, $c_2$ and $c_3$.

### The dwell time statistics with only one bound state

If there is only one bound state or equivalently $k_\mathrm{t} = 0$, for a stator unit that first becomes bound at time $t = 0$, its survival probability is given by: $S(t) = \exp(-k_{\mathrm{off,1}}t)$. From the survival probability distribution, we can obtain the dwell time distribution $P(\tau) = -\frac{dS}{dt}|_{t=\tau} = k_{\mathrm{off,1}}\exp(-k_{\mathrm{off,1}}\tau)$, which follows a simple exponential decay function with a single time scale $k_{\mathrm{off,1}}^{-1}$ given by the off rate $k_{\mathrm{off,1}}$. The exponential form of $P(\tau)$ with a single time scale does not agree with the experiments (**Fig. 1** in the main text). The disagreement can also be seen by computing the normalized variance magnitude $V = \frac{\int_0^\infty \tau^2 P(\tau)d\tau}{(\int_0^\infty \tau P(\tau)d\tau)^2} - 1 = 1$ whereas $V \approx 2$ was observed in the experiments as shown in **Fig. 3c** in the main text. Furthermore, with only one bound state, the effective off rate $k_-(N) = Nk_{\mathrm{off,1}}$ becomes independent of time. As a result, we would have $P_\pm(\tau|N) = k_\pm(N)\exp[-(k_-(N)+k_+(N))\tau]$ and the average dwell times for the '+' (on) and the '-' (off) transitions being equal: $\tau_+(N) = \tau_-(N) = (k_-(N)+k_+(N))^{-1}$, which is again inconsistent with experiments, which show that $\tau_+(N) \neq \tau_-(N)$ (see **Fig.** S3). These experimental results thus rule out the simple two state model (one bound and one unbound) for stators.

### The model parameters are obtained with an optimization algorithm

Given that we can derive analytical expressions for the relevant quantities using the model equations, we implemented a gradient descent algorithm to find the rate parameters leading to the best agreement between the theory and the experimental results. More precisely, we tried to find the values of $k_+(N)$, $c_i$ ($i = 1,2,3$), $\sigma_+$ and $\sigma_-$, as defined in the previous sections, that would lead to values of $\langle\tau\rangle$, $f_+$ and $\langle V\rangle_N = \left\langle \sigma_\tau^2/\langle\tau\rangle^2\right\rangle_N$ where $\langle ...\rangle_N$ indicates an average over $N$, that are closest to what can be directly obtained from experiments. To do this, we define a vector parameter $p$, whose components are the parameters that we are trying to fit, and a loss function $L$ defined as

$$L(p) = \sum_{N=0}^{9}\left[\alpha_1(\langle\tau\rangle^{exp} - \langle\tau\rangle^{mod}(p,N))^2 + \alpha_2(f_+^{exp} - f_+^{mod}(p,N))^2\right] + \alpha_3\left(\frac{1}{9}\sum_{N=0}^{9}\sigma_\tau^2/\langle\tau\rangle^2 - V^{mod}\right)^2\,, \tag{S23}$$

where $\alpha_1$, $\alpha_2$ and $\alpha_3$ are three constant weights. At every step the parameter vector is updated according to

$$p \to p - \eta\cdot\nabla_p L\,, \tag{S24}$$

where $\eta$ is a constant learning rate. Here, with *exp* and *mod* we indicate the mean values of each quantity as measured from the experimental data or as obtained from the model equations respectively.

The choices of the parameters $\alpha_i$ ($i = 1,2,3$) depend on the accuracy that we want to have in the fitting of the three experimental variables. In our case we chose $\alpha_1 = 1/\sigma_\tau^2$, where $\sigma_\tau^2 = \left\langle \tau^2 \right\rangle - \langle \tau \rangle^2$, $\alpha_2 = 1$ and $\alpha_3 = 10$.

The fit of our model to the experimental data shown in **Fig. 3** in the main text led to the parameters $c_1 = 0.30 \pm 0.11$, $c_2 = 0.12 \pm 0.14$, $c_3 = 0.06 \pm 0.16$, $\sigma_+ = 0.19 \pm 0.1$, $\sigma_- \leq 0.0005$. The errors correspond to the fluctuation in the value of the parameter that would lead to a 50% increase in the loss function.
